# HIV-specific CD8 T cells from elite controllers have an epigenetic imprint that preserves effector functions

**DOI:** 10.1101/2021.09.28.459512

**Authors:** Adolfo B. Frias, Rachel L. Rutishauser, Ashish A. Sharma, Tian Mi, Hossam Abdelsamed, Caitlin Zebley, Christian M. Constantz, Mars Stone, Michael P. Busch, Steven G. Deeks, Rafick-Pierre Sékaly, Ben Youngblood

## Abstract

Several lines of evidence support a central role for CD8 T cells as key determinants in the control of HIV, particularly in rare “elite controllers” who control the virus to undetectable levels in the blood in the absence of antiretroviral therapy (ART). While HIV-specific CD8 T cells isolated from elite controllers have enhanced antiviral cytokine production and proliferative capacity in response to antigen stimulation when compared to cells isolated from viremic or even aviremic ART-suppressed non-controllers, the cell-intrinsic mechanisms underlying the enhanced T cell memory-like function of HIV-specific CD8 T cells in elite controllers remain largely undefined. To identify the transcriptional and epigenetic pathways that regulate functional capacity in HIV-specific CD8 T cells in elite controllers, we performed genome-wide transcriptional and DNA methylation analysis of MHC Class I multimer+ CD8 T cells sorted from aviremic elite controllers compared to aviremic non-controllers on suppressive ART. Co-omics analysis revealed enrichment for gene signatures that support a multipotent differentiation state, cell survival, and a long-lived effector cell fate in HIV-specific CD8 T cells from elite controllers. Specifically, we observed DNA methylation programs at the transcription factor binding sites of the stem-associated factors TCF-1 and LEF1 that delineate HIV-specific CD8 T cells from elite controllers versus ART-treated individuals. HIV-specific CD8 T cells in elite controllers also maintain T cell receptor and IL-12/STAT4 pathway signaling and have suppressed pro-apoptotic TNFα pathway signaling. These findings show that HIV-specific CD8 T cells from elite controllers have enhanced expression and DNA methylation programs that maintain developmental potential and in turn promote long-term survival, proliferative potential, and effector capacity. These data also provide new insights into the relationship between stem-associated transcription factors and stable epigenetic restriction of T cell developmental capacity.

## Introduction

The majority of people with Human Immunodeficiency Virus (HIV) infection initially generate a robust immune response but ultimately fail to control of the virus, resulting in persistent infection and the subsequent development of acquired immunodeficiency syndrome (AIDS). A rare group of individuals (<1% of people with HIV), known as elite controllers (ECs), control the virus to undetectable levels in the blood and do not progress to AIDS. Several lines of evidence, including genome-wide association studies in humans and experimental studies in macaques with Simian Immunodeficiency Virus (SIV) support the notion that CD8 T cells play an essential role in mediating viral control in elite controllers (1–3).

HIV-specific CD8 T cells isolated from ECs produce more cytokines and proliferate more robustly in response to T cell receptor (TCR) stimulation than their counterparts in untreated non-controllers with high viral loads (4–7). Interestingly, HIV-specific CD8 T cell functional capacity in non-controllers is not completely restored by suppressive antiretroviral therapy (ART) (8,9), suggesting that persistent exhaustion and impaired responsiveness to antigen stimulation may be epigenetically regulated in these cells, as has been seen in other settings (e.g., after virologic cure in chronic hepatitis C infection, (10–13). Based on murine models of infection with lymphocytic choriomeningitis virus (LCMV), compared to functional effector and memory cells generated in the context of an acute infection, exhausted virus-specific CD8 T cells occupy a unique transcriptional, epigenetic, and metabolic state imprinted early during chronic infection (14–16). In particular, in both mice and humans, the transcription factor (TF) Tox (17), *de novo* DNA methylation by the methyltransferase DNMT3a (18) (Prinzing et al., *Sci Transl Med*, accepted), and the preferential use of glycolytic bioenergetic pathways (16) have been shown to promote CD8 T cell exhaustion in humans. In HIV infection, exhausted HIV-specific CD8 T cells isolated from viremic non-controllers express higher levels of co-inhibitory receptors that inhibit TCR signaling (e.g., PD-1 (19)) and lower levels of the T cell stemness and memory-associated TF TCF-1 (encoded by the gene, *TCF7*) compared to the more functional HIV-specific CD8 T cells in ECs (9,20). While chromatin accessibility differences persist in non-controllers after HIV viral load is suppressed with ART (10), the transcriptional and underlying epigenetic mechanisms that regulate persistent exhaustion in this setting have not been elucidated, nor is it clear what mechanisms promote enhanced functional capacity in HIV-specific CD8 T cells in ECs. Because many HIV cure strategies such as therapeutic vaccination and adoptive T cell therapies seek to promote the formation of EC-like functional HIV-specific CD8 T cells in ART-suppressed individuals, it is critical to understand how exhaustion and virus-specific CD8 T cell functional capacity is regulated in these two clinical groups.

In this study, we compared the transcriptional and epigenetic profiles of MHC Class I multimer+ HIV-specific CD8 T cells from aviremic ECs versus aviremic ART-suppressed non-controllers to understand why HIV-specific CD8 T cells from ECs maintain long-term enhanced responsiveness to TCR simulation. Using whole-genome RNA-sequencing and DNA methylation profiling, we found that genes associated with distinct T cell differentiation states (e.g., effector, memory/stem-like, exhausted) were differentially expressed and had regions of differential DNA methylation in HIV-specific CD8 T cells from ECs compared to non-controllers on suppressive ART. The retained memory/stem-like functional capacity of EC virus-specific T cells was coupled to distinct DNA methylation programs at TCF-1 and LEF1 transcription factor binding sites. Furthermore, functional HIV-specific CD8 T cells from ECs have transcriptomic and DNA methylation signatures suggestive of a more active metabolic state and heightened responsiveness to TCR, IL-12/STAT4, IL-2/STAT5 and GPCR signaling. This study provides insight into transcriptional and epigenetic mechanisms that govern sustained proliferative capacity, survival, and antiviral function of HIV-specific CD8 T cells in elite controllers and identifies candidate target pathways for optimizing T cell-based HIV cure therapies.

## Results

### HIV-specific CD8 T cells from ECs versus ART-suppressed non-controllers are transcriptionally and epigenetically distinct

During HIV infection, CD8 T cells from aviremic ECs have enhanced proliferative capacity and cytokine production compared to cells from aviremic non-controllers on suppressive ART (8,9). To understand the cell intrinsic mechanisms regulating exhaustion in HIV-specific CD8 T cells from non-controllers on ART compared to ECs, we took a co-omics approach to probe gene expression and epigenetic differences. We FACS-sorted HIV-specific MHC Class I multimer+ CD8 T cells from ECs and non-controllers on suppressive ART enrolled in the San Francisco-based SCOPE cohort and performed bulk RNA sequencing and whole genome DNA bisulfite sequencing. To understand the relationship between the transcriptional signature of HIV-specific multimer+ CD8 T cells and bulk CD8 T cell subsets, we performed a principal component analysis (PCA). Bulk CD8 T cells sorted from the same EC and ART donors using markers that denote classically-defined subsets (naïve [TN: CD45RA+CCR7+CD27+], central memory [TCM: CD45RA+CCR7+CD27+], and effector memory [TEM: CD45RA+CCR7+CD27-]) grouped according to developmental state. As expected, based on their phenotype (9), HIV-specific multimer+ CD8 T cells from both the EC and ART group clustered closely to TEM CD8 T cells. (Fig. 1a). There were 1002 differentially expressed genes (DEGs) between the EC and ART HIV-specific CD8 T cells (p < 0.05; Fig. 1b). Notably, genes more highly expressed in HIV-specific CD8 T cells from ECs suggest enhanced responsiveness to cytokine signaling (e.g., higher expression of *IL2RA*, the gene encoding CD25), faster antiviral responses upon TCR engagement (e.g., higher expression of *IFNG* transcript), and enhanced homing to B cell follicles (e.g., higher expression of *CXCR5*; Fig. 1c, top). In contrast to the memory-associated programming observed in the EC HIV-specific CD8 T cells, genes more highly expressed in HIV-specific CD8 T cells from ART-suppressed individuals suggest a cell type that is more terminally differentiated, both phenotypically (with higher levels of expression of NK cell receptors that tend to mark terminal effector cells, e.g., *KLRC4*) and functionally (with expression of cytotoxic molecules, e.g., *GNLY*) (Fig. 1c, bottom). From the transcriptional data we conclude that, although EC and ART HIV-specific CD8 T cells both cluster with the bulk TEM subset, HIV specific CD8 T cells from ECs differentially express transcripts that suggest enhanced T cell survival and homing, and a less differentiated T cell state.

**Figure 1.**
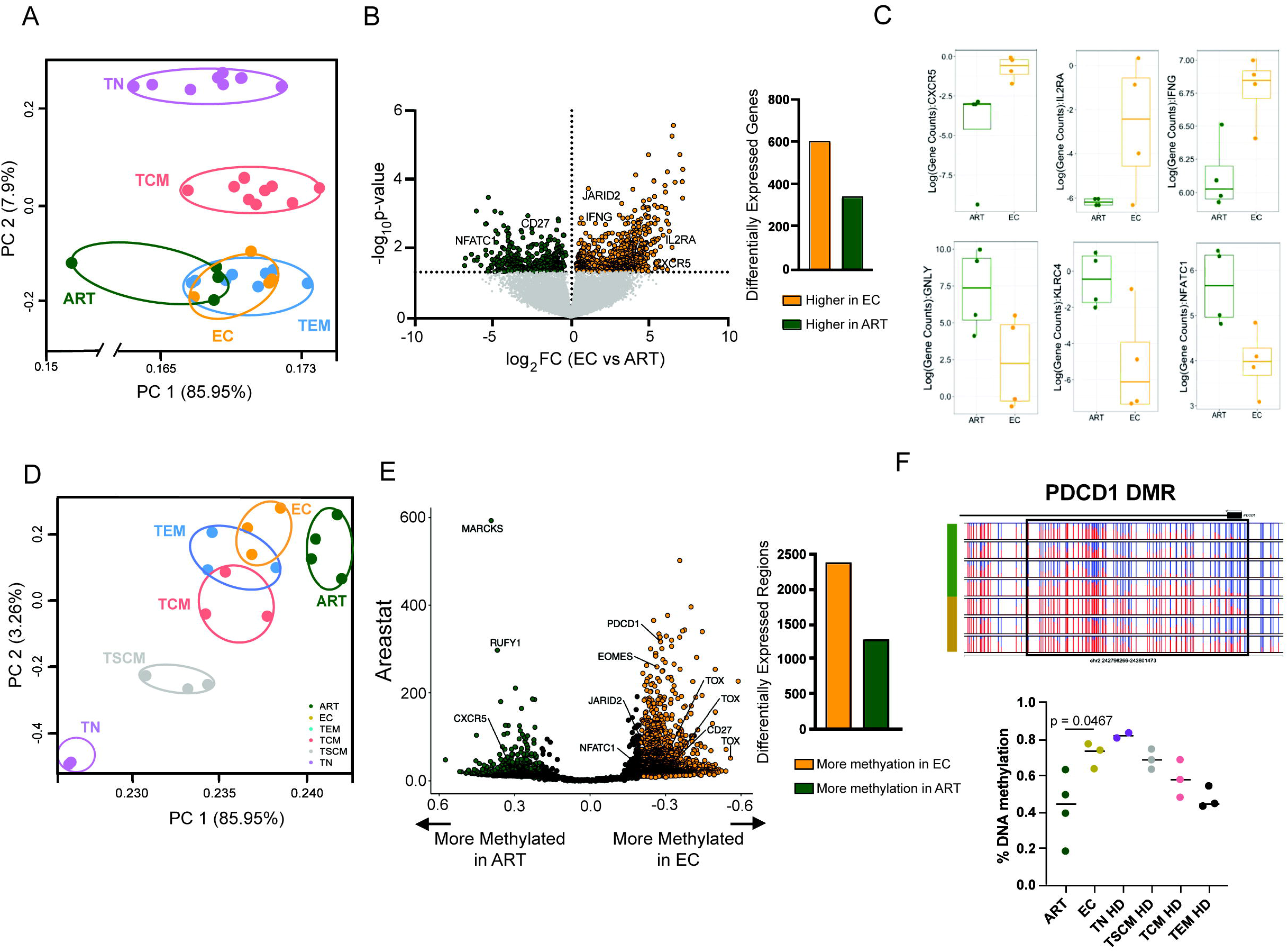
Transcriptional and DNA methylation programs delineate ART versus EC HIV-specific CD8 T cells. A. Principal component analysis (PCA) of RNAseq profiles of sorted HIV-specific multimer+ CD8 T cells from elite controllers (EC) versus ART-suppressed non-controllers (ART) and bulk TN, TCM, and TEM CD8 T cells from the same donors. B. Volcano plot of differentially expressed genes between EC (n = 4) and ART (n = 4) HIV-specific CD8 T cells (p<0.05) C. Representative transcripts enriched in EC (top) and ART (bottom) HIV-specific CD8 T cells D. PCA of DNA methylation profiles of sorted HIV-specific multimer+ CD8 T cells (EC versus ART) and bulk TN, TSCM, TCM, and TEM CD8 T cells from HIV-uninfected donors. E. Genes associated with differentially methylated regions (DMR)s between ART (green) and EC (yellow) HIV-specific CD8 T cells (methylation difference between the two conditions versus area-statistic). DMRs below the 20% threshold are in black. F. DNA methylation patterns at intron 1 of the *PDCD1* locus in HIV-specific CD8 T cells from EC (yellow) versus ART (green) donors (top) and percent DNA methylation in this region in bulk CD8 T cell subsets compared to EC or ART HIV-specific CD8 T cells (bottom). P-value calculated using average methylation from significant DMR CpG sites.

We next performed a PCA analysis on the samples in the DNA methylation dataset. Once again, bulk CD8 T cell populations (TN, TCM, TEM, and stem-cell memory [TSCM: CD45RA+CCR7+CD95+] CD8 T cells isolated from HIV-uninfected donors) clustered by developmental state. While the HIV-specific CD8 T cells from ECs grouped near bulk CD8 TEM cells as in the transcriptome dataset, the ART HIV-specific CD8 T cells clustered separately (Fig. 1d). In order to identify the DNA methylation programs that delineate HIV-specific multimer+ CD8 T cells from ECs compared to ART-suppressed individuals, we performed a pairwise comparison of gene-associated differentially methylated regions (DMRs), defined as significantly differential methylation (p < 0.01) at ≥ 3 CpG sites within a minimum base pair length of 100, and we associated each DMR to a single gene (Fig. 1d). Applying a threshold of 20% methylation difference, we found 3690 DMRs between EC versus ART HIV-specific multimer+ CD8 T cells with the majority of regions (64.9%) more methylated in ECs, and largely located within intron or intergenic regions. Of the DMRs, we found that HIV-specific CD8 T cells from ECs maintained methylation of the first intronic region within the *PDCD1* locus (encodes PD-1), with a methylation pattern at this site similar to more multipotent bulk CD8 T cell subsets (e.g., TN, TSCM; Fig. 1f). Taken together, these data show that HIV-specific CD8 T cells from ECs versus ART-suppressed individuals can be distinguished by their DNA methylation profile. Notably, we found methylation differences in the gene locus encoding the inhibitory receptor PD-1, which suggests one potential mechanism for sustained enhanced TCR responsiveness in HIV-specific CD8 T cells in ECs.

### HIV-specific CD8 T cells from ECs have transcriptional and epigenetic features of long-lived memory cells

To better define the gene expression and/or methylation signatures associated with HIV-specific CD8 T cells isolated from either ECs or ART-suppressed previous non-controllers, we performed gene set enrichment analysis on our transcriptome and methylome datasets using publicly-available gene lists generated from bulk CD8 T cells in different differentiation states (e.g., TN, TSCM, TCM, TEM and the recently defined long-lived effector cells [LLECs: CD45RO+KLRG1+CD127^Int^CD27^low^] (21)), as well as functionally-distinct sub-populations of antigen-specific CD8 T cells in chronic viral infection (e.g., terminally exhausted cells [Tex], progenitor [Prog1, Prog2] and intermediately differentiated sub-populations (22)) (Figure 2a, top). Compared to ECs, multimer+ HIV-specific CD8 T cells from ART-treated individuals had both increased expression and lower DNA methylation of genes that define TEM cells (compared to the less-differentiated TCM and TSCM subsets), and they also had higher expression and lower DNA methylation of genes associated with more terminally exhausted virus-specific CD8 T cells compared to the progenitor or intermediate sub-populations that form in chronic infection in mice. Leading edge genes from these comparisons whose expression was enriched in the ART CD8 T cells included targets involved in CD8 T cell dysfunction *(TNFRSF1B)* (23), terminal effector function (*GZMB*, *GZMM*) (24), and terminal effector differentiation (*IKZF2*, *ZEB2*) (25,26) (Fig. 2b). In contrast, leading edge genes in EC HIV specific CD8 T cells included markers for lymph node trafficking (SELL), positive modulation of T cell effector metabolism (MYC) (27), and follicular trafficking and CD8 T cell mediated viral control (CXCR5) (28) (Fig 2a, top). Similarly, genes associated with DMRs that were less methylated in the ART samples include the lymph node and bone marrow egress marker *S1PR5* and the T cell effector differentiation marker *CX3CR1* (29) (Fig. 2c). Interestingly, compared to ECs, HIV-specific CD8 T cells from ART-suppressed individuals also had higher expression of genes that are more highly expressed in TCMs compared to LLECs, a recently-described population of cells that are long-lived but simultaneously maintain immediate effector functions (21). Taken together, our data suggest that HIV-specific CD8 T cells from ECs appear to occupy a less exhausted hybrid differentiation state in which they retain some features of more multipotent memory cells as well as some features of long-lived effector cells.

**Figure 2.**
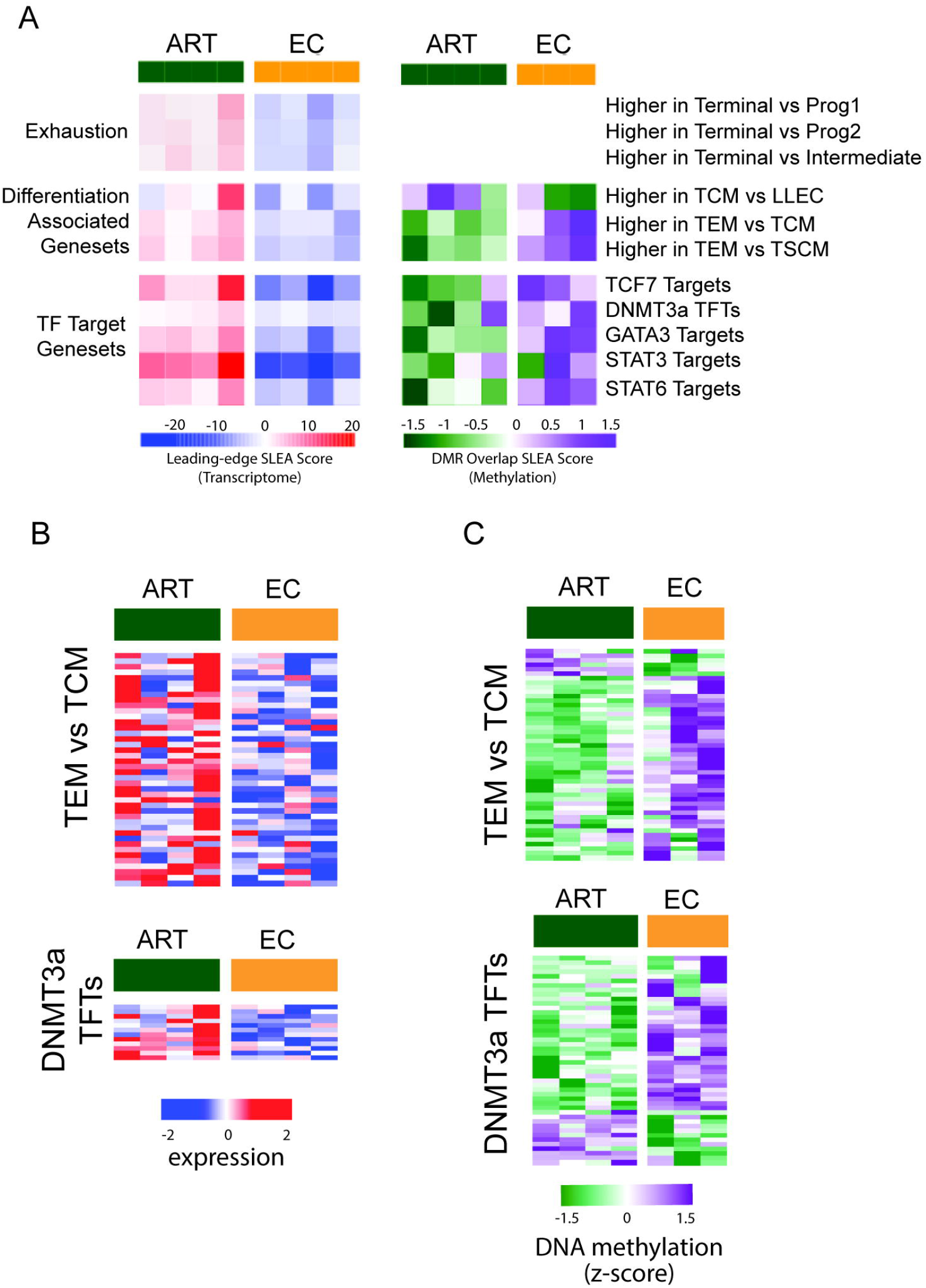
Pathways that govern CD8+ T cell memory and exhaustion are differentially regulated in HIV-specific CD8+ T cells from elite controllers versus ART-suppressed non-controllers. A. Heatmap of pathways that are differentially expressed and methylated between EC and ART HIV specific CD8 T cells (Nominal p value < 0.01). B. Heatmap of leading edge genes from pathways differentially expressed and methylated. C. Heatmap of DMRs from pathways differentially expressed and methylated

### Distinct regulation of T cell differentiation transcriptional and epigenetic pathways in HIV-specific CD8 T cells from ECs

Our prior work on the DNA methyltransfyrase, DNMT3a, has found that DNMT3a-mediated *de novo* DNA methylation reinforces the terminal exhaustion state of mouse and human T cells that have been chronically stimulated (18,30). Comparing the EC versus ART HIV-specific CD8 T cell DMR list with that of our established DNMT3a gene signature, we found that more than 20% of the DMRs overlapped with our DNMT3a gene signature. Furthermore, when we assessed the expression of transcripts that encode DNMT3a-targeted transcription factors, we found that they were expressed at significantly higher levels in ART compared to EC HIV-specific CD8 T cells (Fig 2XX). This pattern suggests increased DNMT3a activity may be one potential mechanism by which exhaustion is reinforced in HIV-specific CD8 T cells from non-controllers.

Because virus-specific CD8 T cell effector/memory differentiation is regulated by the coordinated activity of several hallmark transcription factors (31), we were interested in the broad differential regulation of transcription factor target genes between EC versus ART HIV-specific CD8 T cells. ART HIV-specific CD8 T cells were enriched for the expression of STAT3 targets (and had lower methylation at these target genes), and they also were enriched for the expression of targets of two type-2 cytokine signaling factors (STAT6 and GATA3), which have been shown to impair CD8 T cell division and differentiation (32) (Fig. 2a bottom). Leading edge genes from the STAT3 targets that were both more highly expressed in ART and also had an associated DMR compared to the EC samples included the exhaustion-associated factors *NFATC1* and *TOX* and the type-2 immunity/NK developmental TF *NFIL3* (data not shown). Other leading-edge genes from the STAT3 target list that were more highly expressed in the ART HIV-specific CD8 T cells suggest terminally differentiated CD8 T cells (*IKZF2*), diminished proliferation and survival (*AKT2*), and DNMT3a activity (*ZBTB18*) (33). On the other hand, STAT3 target genes associated with DMRs that demonstrated a trend towards higher expression in HIV-specific CD8 T cells from ECs included a guanine nucleotide exchange factor downstream of TCR and CD28 signaling (*VAV3*), a receptor component for the cytokine IL-12 (*IL12RB1*), and a stem-associated transcription factor (*BACH2*; data not shown). Lastly, from the STAT3 targets we also observed differential methylation of the B cell follicle homing molecule *CXCR5* (p = 0.043; also a differentially expressed gene, Fig. 1c). These data show that, compared to ECs, CD8 T cells isolated from ART-suppressed individuals have differential DNA methylation at and are enriched for the expression of genes associated with type-2 immune signaling, terminal differentiation, and exhaustion, suggesting additional potential mechanisms that may underlie their relatively impaired ability to signal downstream of TCR and cytokine stimulation.

To further resolve the mechanisms involved in preserving the functional capacity of HIV-specific CD8 T cells from ECs, we performed Hypergeometric optimization of motif enrichment (HOMER) analysis (34) to identify transcription factors (TFs) whose binding site motifs were over-represented in the DMRs between EC versus ART HIV-specific CD8 T cells. As noted above, the majority (64%) of our DMRs were located within intronic or intergenic regions. Amongst regions that were less methylated in EC (versus ART) HIV-specific CD8 T cells, we noted a striking pattern of enrichment for the binding site sequences for several stem-associated TFs (e.g., HOX9, TCF-1, OCT4-SOX2-TCF-NANOG-POU, LEF1; Fig. 3a). Among the genes associated with the 242 DMRs containing TCF-1 motifs, we identified targets that were less methylated in ECs such as the type-2 immune TF *NFIL3* (intergenic) and the JUN and FOS transcript repressor *PRDM11* (intron), and targets that were less methylated in the ART-suppressed samples such as the receptor tyrosine kinase antagonist, *SPRY2* (35) (intergenic) (Fig. 3b). Interestingly, this striking difference in methylation at TCF-1 motifs between EC versus ART HIV-specific CD8 T cells was not accompanied by differences in the level of expression of the *TCF7* transcript itself; although we have observed differences at the protein level (9).

**Figure 3.**
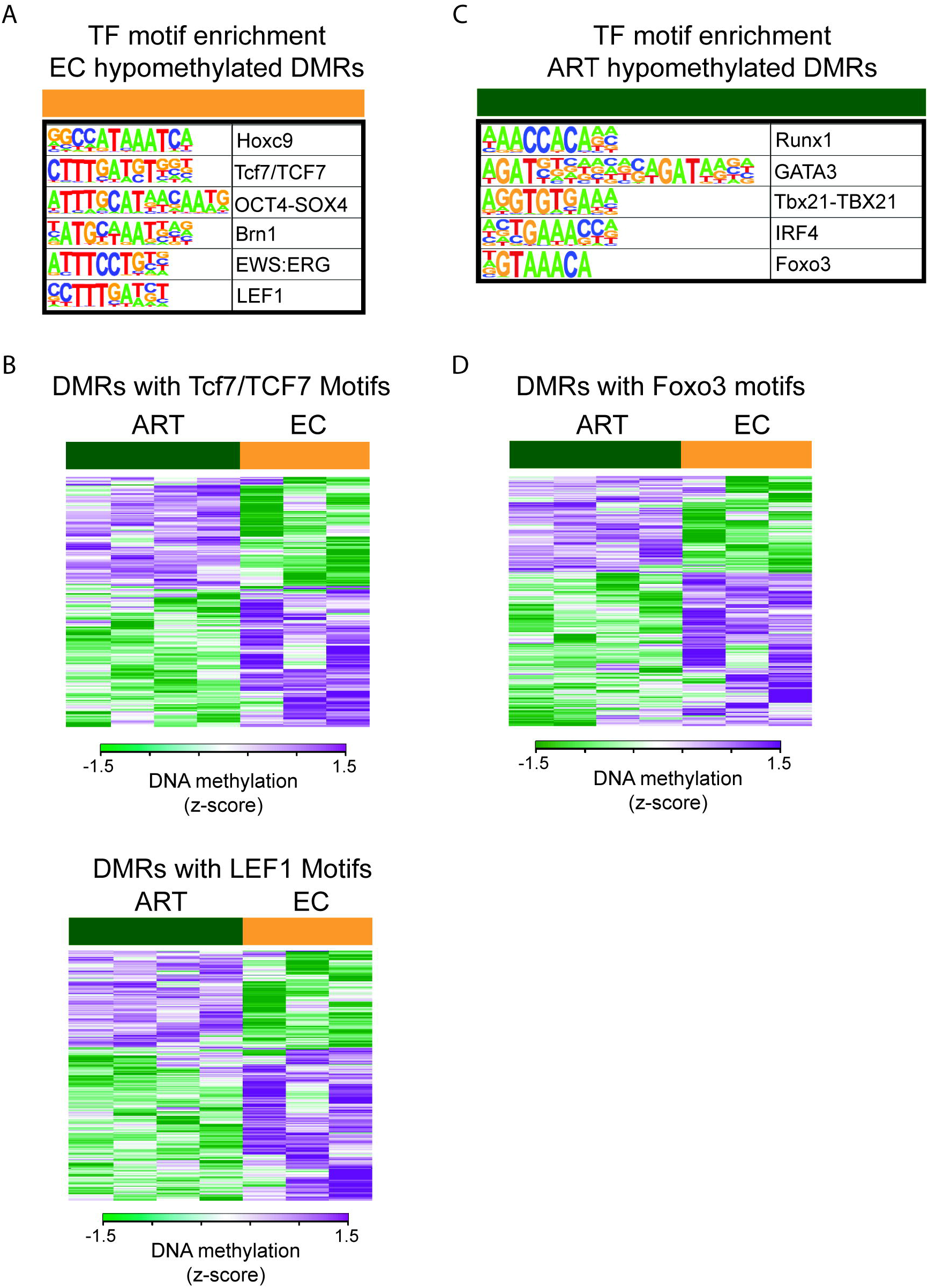
Differential methylated regions at stem-associated transcription factor binding sites delineate ART vs EC HIV specific CD8 T cells. A. HOMER analysis of relatively hypomethylated EC HIV specific CD8 T cell DMRs (FDR < 0.01) B. Heatmap of DMR genes containing Tcf7 motifs C. HOMER analysis of hypomethylated ART HIV specific CD8 T cell DMRs (FDR < 0.01) D. Heatmap of DMR genes containing Foxo3 motifs

In contrast to the ECs, DMRs that were less methylated in ART HIV-specific CD8 T cells were enriched for the binding site sequences for several effector differentiation-associated TFs (e.g., TBX21, IRF4) as well as for FOXO3 motifs (Fig. 3c). Of note, *FOXO3* itself was differentially methylated and was also among the leading-edge genes from the STAT3 and DNMT3a target lists (Fig. 2a). Hypomethylated DMRs in ART HIV-specific CD8 T cells included markers associated with a terminal/TEM fate (*S1PR5* (36))_or CD8 T cell exhaustion (*TOX*, *CD101* (37)). HOMER analysis of the DMRs found enrichment of TCF-1 motifs in DMRs that were relatively hypomethylated in HIV-specific CD8 T cells from ECs and enrichment of FOXO3 motifs in DMRs that were relatively hypomethylated in ART HIV-specific CD8 T cells. In addition, FOXO3 targets, could potentially limit cell cycling and prevent homeostatic proliferation required for survival of long-lasting memory T cells

### Gene expression pathways downstream of TCR and cytokine signaling are differentially regulated in HIV-specific CD8+ T cells from ECs

Given that enhanced functional capacity of HIV-specific CD8 T cells in ECs is measured by increased responses to antigen stimulation (as measured by either cytokine production and/or proliferation) (9), we hypothesized that HIV-specific CD8 T cells from ECs might be transcriptionally and/or epigenetically poised to respond to TCR signals. Indeed, leading edge genes from several of the pathways that we found are enriched in EC HIV-specific CD8 T cells included several genes that encode key proteins involved in TCR signaling (e.g., *ITK*, *LCK*, *SLP76/LCP2*) (Fig. 4c). Furthermore, HIV-specific CD8 T cells also had evidence of increased signaling downstream of IL-12/STAT4 (Fig. 4a,c), a cytokine that is critical for promoting the acquisition of antiviral properties in T cells. Leading edge genes downstream of these pathways that were enriched in the EC HIV-specific CD8 T cells (e.g., *IL12RB1*, *IL12RB2*, *TYK2*) suggest that these cells may have enhance responsiveness to IL-12. Finally, HIV-specific CD8 T cells from ECs also had higher expression of gene targets of pathways that promote metabolic activity and cell cycling (e.g., pathways regulated by MTOR, MYC, and E2F) (data not shown). Taken together, these data show that EC CD8 T cells maintain TCR and IL-12/STAT4 signaling capacity along with enrichment of metabolic pathways important for proliferation (MYC and E2F), effector function (Fatty Acid Metabolism and MTORC1 signaling), and survival (unfolded protein response [UPR]).

**Figure 4.**
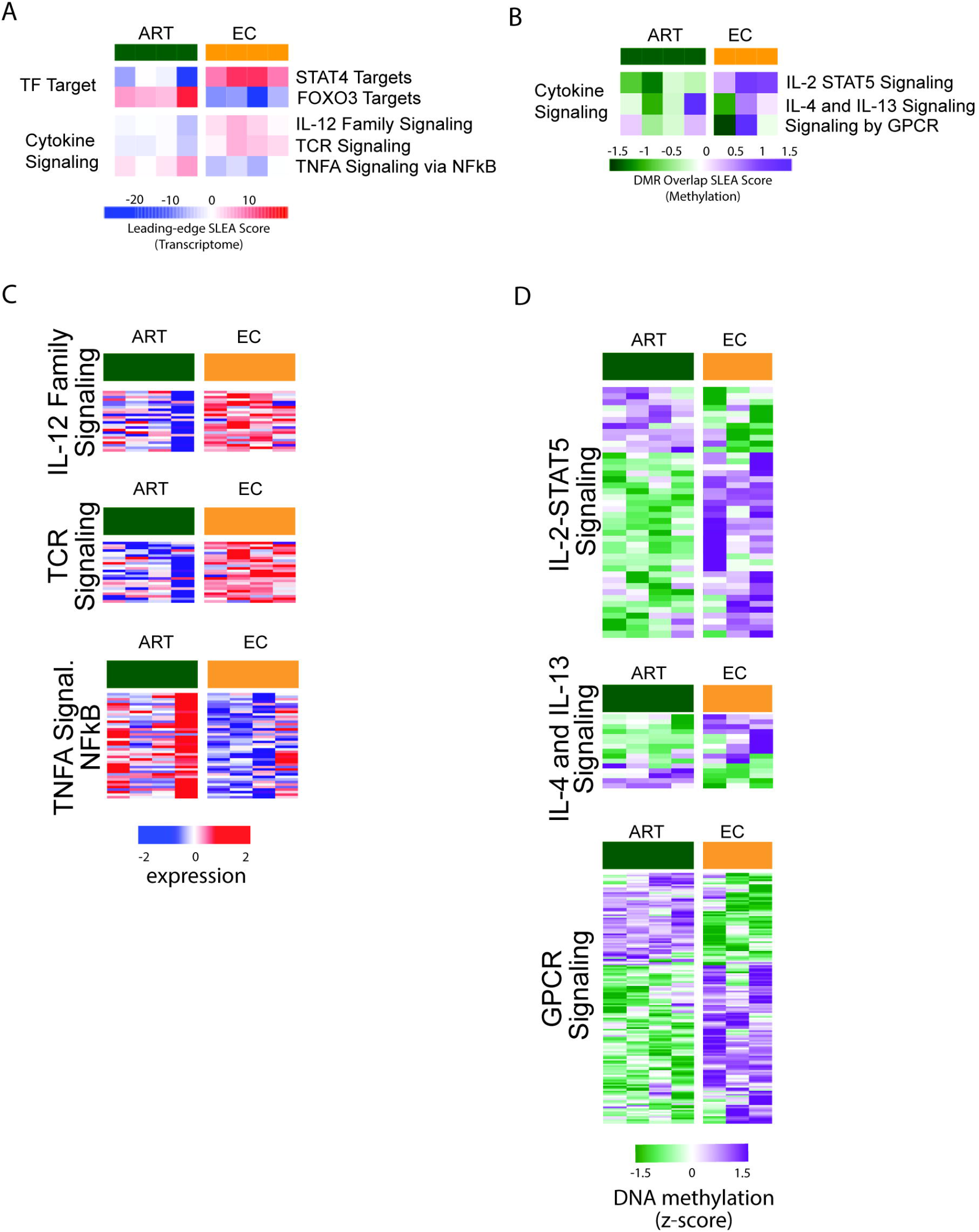
Elite controller HIV-specific CD8+ T cells differentially regulate pathways sensitivity to TCR and cytokine signaling and metabolism. A. Heatmaps of differentially expressed pathways between EC and ART HIV-tetramer+ CD8 T cells (Nominal p value < 0.01). B. Heatmaps of differentially methylated pathways between EC and ART HIV-tetramer+ CD8 T cells. C. Heatmaps of pathway leading edge genes expressed between EC and ART HIV-tetramer+ CD8 T cells. D. Heatmaps of pathway targets differentially methylated between EC and ART HIV-tetramer+ CD8 T cells.

In contrast to ECs, ART HIV-specific CD8 T cells were enriched for TNFα Signaling via NFKB pathway (Fig. 4a). Leading edge genes expressed at higher levels in ART CD8 T cells included the NFKB signaling inhibitors *NFkBIA* and *TNFAIP3* (a.k.a *A20*), and the pro-apoptotic molecules *KLF4* and *DRAM1*. These data suggest that HIV-specific CD8 T cells from indiviudals on ART may have enhanced sensitivity to the negative effects of TNFa including reduced signaling through NFkB and increased pro-apoptotic markers.

Other pathways with gene targets that were differentially methylated between EC and ART HIV-specific CD8 T cells included the IL2-STAT5 signaling, IL-4 and IL-13 signaling, and signaling by G protein-coupled receptor (GPCR) pathways (Fig. 4d). Many differentially methylated genes within these signaling pathways suggest enhanced responsiveness of EC HIV-specific CD8 T cells to TCR and cytokine signaling, including genes that encode isoforms of diacylglycerol (DAG) kinases (e.g., *DGKB*, *DGKK*, *DGKH*, *DKGZ*) and protein kinase C (e.g., *PRKCA*, *PRKCD*, *PRKCH*), guanine nucleotide exchange factors (GEFs; e.g., *SWAP70*, *ARHGEF3*, *VAV3*), the solute transporter *SLC2A3/GLUT3*, GTPase *RHOB*, T cell differentiation TFs *IRF4* and *EOMES*, migratory molecules associated with LN egress (*S1PR5*), B cell follicle homing (*CXCR5*), recruitment to inflammatory sites (*CCR2*), and terminal differentiation (*CX3CR1*). These data demonstrate that HIV-specific CD8 T cells from ECs are transcriptionally and epigenetically poised for rapid TCR and cytokine signaling pathway activation, and they have unique epigenetic programs that regulate the expression of migratory molecules associated with T cell function and developmental status.

## Discussion

HIV-specific CD8 T cells in elite controllers can maintain suppressive antiviral activity for decades in infected individuals (38). Although many HIV cure strategies seek to elicit T cells that possess the functional capacity of HIV-specific CD8 T cells from ECs, the cell-intrinsic mechanisms responsible for their ability to sustain long term effector and proliferative potential are not well understood. By analyzing the transcriptome and methylome datasets from antigen-specificity matched CD8 T cells isolated from ECs and aviremic ART-suppressed non-controllers, our work here identified differentially regulated gene expression and epigenetic pathways involved in memory differentiation and exhaustion, proliferation and survival, and TCR and cytokine signaling that delineate a virus specific T cell response among EC versus aviremic ART treated non-controllers.

Consistent with the known antiviral capacity and enhanced antigen responsiveness of HIV-specific CD8 T cells in ECs (5,7,9,39,40), our analysis identified several genes significantly upregulated in EC CD8 T cells involved in T cell survival (*IL2RA*), homing potential to the major HIV reservoirs (*CXCR5*), and effector function (*IFNG*). These data suggest that retained function of CD8 T cells in ECs allow for surveillance and control during viral latency and potential reactivation (41,42). In line with the idea of sustained viral control, our key findings from the epigenetic analysis included DMRs in the exhaustion-associated loci of *PDCD1* and *TOX* that may act as a potential mechanisms to explain how EC HIV specific CD8 T cells remain poised for TCR stimulation despite their long-term chronic infection setting.

From our combined expression and epigenetic pathway analysis, we observed that EC CD8 T cells were enriched for the expression of genes associated with less exhausted/more multipotent T cell differentiation states (e.g., TSCM, TCM, Tpex) as well as a signature for long-lived effector cells (21) relative to their ART counterparts. Related to sustained long term effector function, our work found expression and DNA methylation pathways that highlighted IL-12 signaling in EC HIV-specific CD8 T cells. Studies have reported the potential of IL-12 to enhance the function of exhausted T cell in tumor and viral infection (43,44) including better tumor control and lower expression of PD-1 (44). This potentially hints at an IL-12 dependent mechanism in EC HIV specific CD8 T cells that is necessary to maintain long term effector CD8 T cell anviral responses while simultaneously retaining stem-like properties.

Previously we showed that the stem-associated TF TCF-1 contributes to viral control differences between EC and ART CD8 T cells (9). Although we did not observe stem-associated TF expression differences between EC and ART HIV specific CD8 T cells in this study, our analysis found differential epigenetic regulation of TCF-1 and LEF1 DNA binding sites that may be responsible for sustained viral control. Our work also found differential regulation of transcription factors targeted by DNMT3a which we have previously found to be responsible for *de novo* DNA methylation mediated exhaustion programs in LCMV chronic mouse model of infection (18). The enrichment of TCF-1 binding site motifs at EC hypomethylated regions suggests an EC HIV specific CD8 T cell intrinsic mechanism that is resistant to DNA methylation at specific sites that enables continued binding of TCF-1. These data suggest that both DNA methylation and demethylation programs may play a role in the establishment of or resistance to DNA methylation-associated exhaustion programs.

In summary, here we have provided a co-omics approach to identify differentially regulated transcriptional programs between HIV-specific CD8 T cells in ECs versus ART-suppressed individuals. Our expression and methylome pathway analysis determined that EC HIV-specific CD8 T cells maintained a more stem-like profile compared to those from ART. We found TCR and cytokine intracellular signaling pathways that remained poised in EC HIV specific CD8 T cells, specifically related to the IL-12 family signaling pathway. In addition, while we didn’t observe direct mRNA expression differences in the stem-associated TFs *TCF7* and *LEF1*, we did find differentially epigenetic regulation of their DNA binding motifs. Future work will focus on whether these findings extend to other HIV-specific epitopes and chronic infection settings. It also remains to be tested as to what DMRs are necessary to phenocopy the sustained antiviral response of the EC HIV specific CD8 T cells. A recent study has shown that it is possible to leverage CRISPR technology to target precise regions for DNA methylation (45) to test potential DMR targets from this study to attempt to copy the EC HIV specific CD8 T cell phenotype. The importance in the characterization of peripheral HIV specific CD8 T cells lies in their immunotherapeutic potential in light of recent findings showing that future immunotherapies may be dependent on recruitable cells considering that tissue resident cells can be refractory to immune checkpoint blockade (46,47).

## Methods

### Human study participants and samples

This study sampled de-identified PBMCs retrospectively collected under IRB approval from participants with HIV enrolled in the Zuckerberg San Francisco General Hospital clinic-based SCOPE cohort.

### Human PBMC sorting by flow cytometry

Cryopreserved PBMCs were thawed as described previously and enriched for CD8 T cells by negative selection with magnetic beads (STEMCELL). HIV-specific CD8+ T cells were identified via staining with peptide–MHC Class I multimers, either monomers that were provided by RPS (Emory University, Atlanta, Georgia, USA) and tetramerized as described using PE, or biotinylated pentamers (ProImmune) followed by staining with streptavidin PE. The multimers used in this study targeted the following specificities: HLA-A*02 (SL9), HLA-B*27 (KK10), HLA-B*07 (TL10). PBMCs were incubated with multimer diluted in PBS for 15 minutes at room temperature (pentamer) or 20 minutes at 37°C (tetramer), followed by 20 minutes at room temperature with surface antibodies (with streptavidin for stains with pentamer), along with a viability dye to allow for discrimination of dead cells (Thermo Fisher Scientific). Live multimer+ and non-multimer bulk CD8 T cell populations were sorted using a BD FACS-Aria II.

### Genomic methylation analysis

DNA was extracted from the sorted cells using a DNA extraction kit (QIAGEN) and then bisulfite-treated using an EZ DNA methylation kit (Zymo Research), which converts all unmethylated cytosines to uracils. WGBS was performed as described previously. Briefly, bisulfite-modified DNA sequencing libraries were generated using the EpiGenome kit (Epicentre) according to the manufacturer’s instructions. Bisulfite-modified DNA libraries were sequenced using Illumina HiSeq 4000 and NovaSeq 6000 systems. Sequencing data were aligned to the Hg19 genome using the BSMAP v. 2.74 software. Using significantly methylated CpG sites (p < 0.01), DMRs were defined as ≥3 CpG sites within a minimum base pair length of 100, and each site associated with a single gene and a pairwise comparison done between EC and ART CD8 T cells.

### Hypergeometric optimization of motif enrichment (HOMER)

A 20% minimum difference in methylation was applied to the EC vs ART HIV-tetramer+ CD8 T cell DMR list and split into DMRs that were relatively hypomethylated in ART or EC CD8 T cells. These DMRs were then analyzed for transcription factor motifs using HOMER. Motifs with an FDR of <0.01 were assessed.

## DECLARATION OF INTEREST

Drs. Youngblood and Zebley have patents related to epigenetic biomarkers and methods for enhancing T cell function for cellular therapies.

## Acknowledgments

The content is solely the responsibility of the authors and does not necessarily represent the official views of the NIH. This work was supported by the National Institutes of Health, K23AI134327 (RLR), R01AI114442 and R01CA237311 (BY), loan repayment program and the National Comprehensive Cancer Network Young Investigator Award (CZ), ASSISI foundation support to BY, and the American Lebanese Syrian Associated Charities (ALSAC to BY). Additional support for SCOPE was provided by the amfAR Institute for HIV Cure Researc (SD and RR).

## Notes

### Competing Interest Statement

Drs. Youngblood and Zebley have patents related to epigenetic biomarkers and methods for enhancing CAR T cell function.

